# Saccadic suppression as a perceptual consequence of efficient sensorimotor estimation

**DOI:** 10.1101/099051

**Authors:** F. Crevecoeur, K. P. Kording

**Affiliations:** Institute of Information and Communication Technologies, Electronics and Applied Mathematics (ICTEAM), Université catholique de Louvain (UCLouvain), Belgium; Institute of Neuroscience (IoNS), UCLouvain; Rehabilitation Institute of Chicago, Northwestern University, Chicago, USA

## Abstract

Humans perform saccadic eye movements two to three times per second. When doing so, the nervous system strongly suppresses sensory feedback for extended periods of time in comparison with the movement time. Why does the brain discard so much visual information? Here we suggest that perceptual suppression may arise from efficient sensorimotor computations, assuming that perception and control are fundamentally linked. More precisely, we show that a Bayesian estimator should reduce the weight of sensory information around the time of saccades, as a result of signal dependent noise and of sensorimotor delays. Such reduction parallels the behavioral suppression occurring prior to and during saccades, and the reduction in neural responses to visual stimuli observed across the visual hierarchy. We suggest that saccadic suppression originates from efficient sensorimotor processing, indicating that the brain shares neural resources for perception and control.

## Introduction

People skillfully combine acquired knowledge, and sensory feedback, a combination that is typically modeled using Bayesian statistics ^1,2^. This framework effectively captures behavior in numerous tasks broadly corresponding to perceptual decision-making ^3–7^, or online movement control ^8–11^. Although perceptual decision-making and sensorimotor control are often considered different phenomena, they cannot really be dissociated in the real world – we need to use the same brain for movement and perception ^12,13^. Perceptual decision-making and sensorimotor behaviors may thus be linked.

A salient case of crosstalk between perception and sensorimotor behavior is *saccadic suppression*: visual acuity is reduced around the time of a saccade. It is often assumed that this mechanism maintains stable perception of our surroundings ^14^. However, the behavioral and neural dynamics of saccadic suppression are difficult to explain if it were purely related to compensating for movement-related effects.

Indeed, previous work has shown that saccadic suppression is controlled centrally, and typically lasts for >100ms even for saccadic movements of ~50ms ^15^. Furthermore, simulating the displacement of the retinal image without a saccade does not elicit similar suppression as during real saccades ^16,17^. As well, the reduction of visual acuity was reported to selectively impact the neural pathways contributing to motion perception (i.e., the magnocellular stream), without complete suppression ^18^, while leaving information with high frequency contents or color unaffected ^19,20^. It is unclear why maintaining perceptual stability would require such a long, powerful, and selective suppression of sensory feedback, if it were purely related to perception, and independent of motor control (Fig. 1, separate resources hypothesis, H1). After all, there is a price to be paid to discard so much sensory information for some 100ms. Thus, saccadic suppression is a complex phenomenon, for which a meaningful function has not been clearly identified.

**Figure 1.**
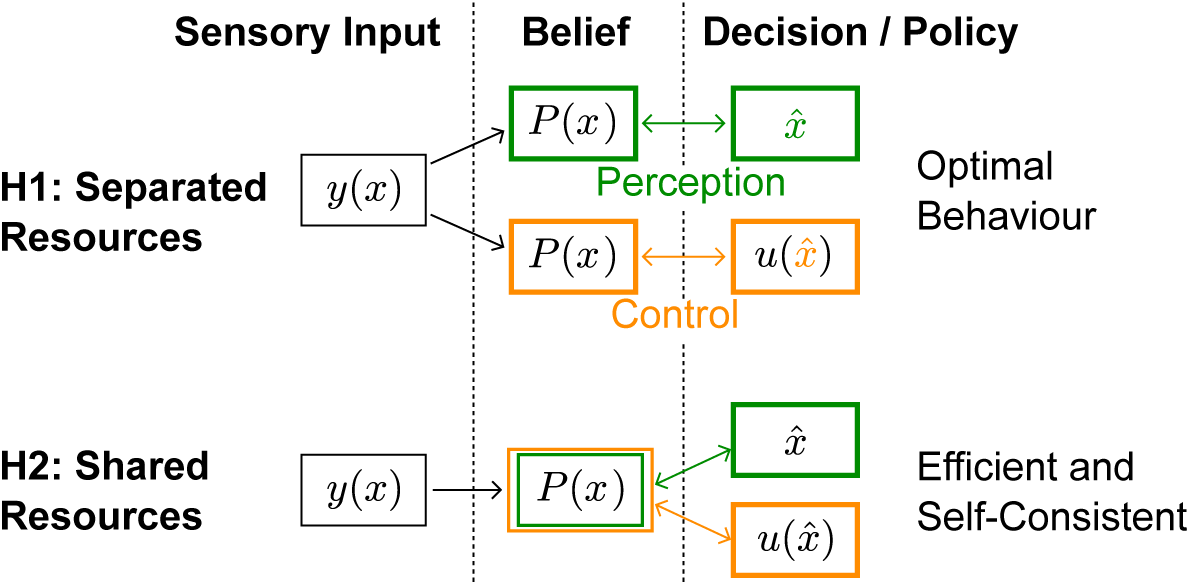
Schematic illustration of two hypotehses. The sensory input, *y* (·) is a function of the variable of interest, *x*. The posterior distribution of *x* is represented by *P*(*x*). The estimate of *x* is designated by 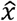, which can be any statistics related to *P*(*x*) such as the posterior mean or mode. Finally the control command is a function of the estimate, represented by 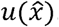 In the hypothesis of separated resources (H1), computations of the posterior belief are carried out independently for perception and control. In this scenario, the uncertainty induced by the control commands does not impact the perceptual estimate. This possible architecture is optimal in the sense that it would minimize loss of sensory information. In the hypothesis of shared resource (H2), the computation of the posterior belief about the state of a variable is shared for perception and control, thus both processes are similarly influenced by control-dependent noise. Although the first hypothesis is optimal, the second hypothesis is more efficient in terms of neural resources, and is also self-consistent (see Discussion).

Here we phrase this problem as a problem of shared resources – the same neural hardware is used for sensorimotor tasks and perceptual decision-making (Fig. 1, shared resources, H2). Formulating this problem in the framework of Bayesian estimation, we show theoretically that several features of saccadic suppression are expected if the brain uses the same posterior beliefs about the state of the eye for perception and control. Our model shows that uncertainty about the instantaneous state of the eye should increase with motor commands as a result of signal-dependent noise and of sensorimotor delays, making delayed sensory information less reliable around the time of movement. In an optimal estimation framework, this gives lower weights to sensory inputs when we move. Our study thus shows how sensorimotor control can give rise to sensory suppression in an efficient brain, provided that neural resources are shared between perception and control.

## Results

### An optimal control model of eye movements

If we want to explore the relationship between saccadic suppression and control we need to model the underlying system. First, the nature of the representation matters: although saccades are often simplistically viewed as ballistic (or *open-loop*) movements, these movements are monitored online through the corollary discharge ^21–25^. Second, sensory feedback matters: we are not “blind” during saccades. There is no peripheral interruption of sensory inflow, and information about specific spatiotemporal frequency or color is still good ^19,26^. Moreover, target jumps during long saccades can influence movement ^27,28^. We should thus model saccades as driven by closed-loop control (Fig. 2a).

**Figure 2.**
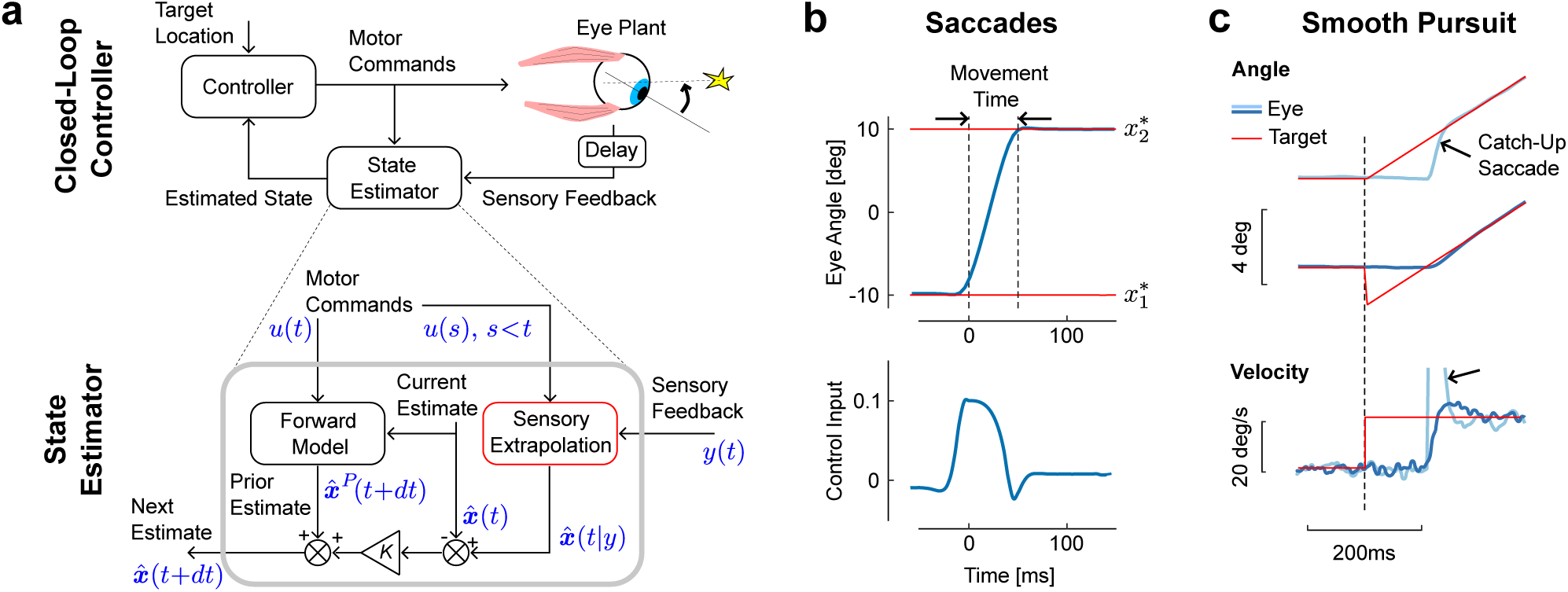
**a.** Schematic representation of the control and estimation architectures. We consider a closed loop controller based on optimal feedback control and state estimation. The dynamics of the eye plant corresponded to a second order system with time constants taken from the literature (13ms and 224ms). Bottom: Optimal state estimator based on usual Kalman filtering, and augmented with the extrapolation of sensory feedback to compensate for sensorimotor delays (Sensory Extrapolation, red box). The symbolic representation of the signals in blue follows the same notations as in the Methods: ***y***(*t*) is the sensory feedback,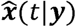 is the extrapolation of sensory feedback, *u*(·) is the sequence of previous and current control commands,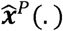 and 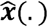 are the prior and posterior estimates at the corresponding time steps. **b.** Top: Modeled saccadic eye movement from the first 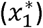 to the second fixation target 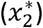. Bottom: Associated control function. Time zero corresponds to the end of the fixation period to the first target, **c.** Illustration of the sensory extrapolation performed in the state estimator. The simulated task is to track the target, which suddenly starts moving (velocity jump) with or without position jump in the opposite direction. The simulated eye trajectory shows how the extrapolation of target motion over the delay interval generates a catch up saccade (black arrow). This compensatory movement is also illustrated in the velocity trace.

To describe saccades in the context of closed loop control, we model a controller that takes the sensory feedback and the corollary discharge as input, and outputs motor commands. We employ a Linear-Quadratic-Gaussian (LQG) controller, which can deal with noise both in sensory feedback and control signals. As model for the oculomotor plant we use a second order model (see Methods). This explicit model of saccadic control allows us to derive predictions of eye movement behaviors and gives us a control process that we can relate to saccadic suppression.

The important feature of this control design in the context of this paper is its state estimator. The control of saccadic eye movements relies on the corollary discharges as well as on sensory feedback, which jointly allow state estimation. This state estimator has two main components. The first is a forward model that dynamically updates the current estimate based on the corollary discharge (Fig. 2a, bottom: Forward Model). The output is a prior estimate of the next state at the next step. The second component is the sensory extrapolation, which combines the delayed sensory feedback with the corollary discharge to estimate the current state (Fig. 2a: Sensory Extrapolation, red). This sensory extrapolation is critical for the behavior of the model.

The presence of sensory extrapolation is supported by previous studies showing that error signals used to generate saccades depend on an estimate of the current state of the eye or of the target ^29–33^, which clearly requires extrapolation of sensory feedback. Indeed, because the system only has access to the delayed feedback, this feedback must be extrapolated prior to compare it with the one-step prediction, or prior. This operation does not appear explicitly in standard control models in which sensorimotor delays were considered^10,11,34,35^, because these previous studies used system augmentation, and the sensory extrapolation in this case falls out of the block-structure of the model. However, this component is necessary, and ignoring it can lead to instability^35^. In the present model, this sensory extrapolation is performed explicitly (Eqn. 5), which also allows incorporating the impact of signal-dependent noise in the extrapolation (see also Methods). The model can then correct the one step prediction, weighting the difference between feedback and expected feedback optimally (Fig. 2a, *K*(*t*) is the Kalman gain).

### Saccades and smooth pursuit

The first behavior that our model must describe is a saccade. The model reproduces stereotyped, step-like trajectory (Fig. 2b, top), like those found during real saccades. Moreover, the associated commands presents with a typical wide agonist burst, followed by short and sharper antagonist activity that stabilizes the eye at the goal target (Fig. 2b, bottom). This pattern of control, shaped by the fast time constants of the oculomotor plant, is compatible with the pattern of burst neurons that generate saccades ^21^. Thus the model replicates both behavioral and physiological aspects of saccadic eye movements.

A second behavior that our model can capture is smooth pursuit. We do not imply that these two behaviors are supported by the same neural hardware, and the model does not make any prediction about their neural implementation. Instead, we simply assume that optimal state estimation underlies both saccades and pursuit, which is in agreement with the hypothesis that these movements are distinct outputs of shared sensorimotor computations ^36,37^. The model reproduces typical responses to changes in target velocity, occurring with or without initial target jump (Fig 2c). When the target starts moving (velocity jump), position error accumulates over the delay interval, which in turn requires a rapid compensatory movement to catch up with the target (Fig. 2b, light blue). Although the controller was not explicitly designed to model the interaction between saccades and pursuit, the catch-up saccade in Fig. 2b naturally falls out of the simultaneous correction for errors both in position and velocity. In contrast, when the target jumps backwards at the onset of the velocity jump (Fig. 2c, dark blue), the eye starts moving smoothly and there is no catch-up saccade ^38^. The model also reproduces corrections following perturbations applied during movement through internal monitoring of the corollary discharge, as well as online corrections for target jumps occurring during long saccades (simulations not shown). In all, the model generates typical trajectories and control commands associated with eye movements, and reproduces the dynamic estimation of the target resulting from the sensory extrapolation.

### Saccadic Suppression as a Consequence of Optimal Estimation

Our muscles produce signal dependent noise; the stronger the muscles pull, the more noisy the state. The phasic activity associated with the agonist burst induces a peak in the variance of the control signal (Fig. 3a, solid). Thus motor commands produce instantaneous noise, and because of the delay, there is no way for the nervous system to directly subtract or filter out this noise. As a consequence, the extrapolation error computed over an interval that includes even a fraction of the control burst has higher variance. In other words, moving the eye effectively induces visual uncertainty, which can only go back to baseline after the end of muscle activation.

**Figure 3.**
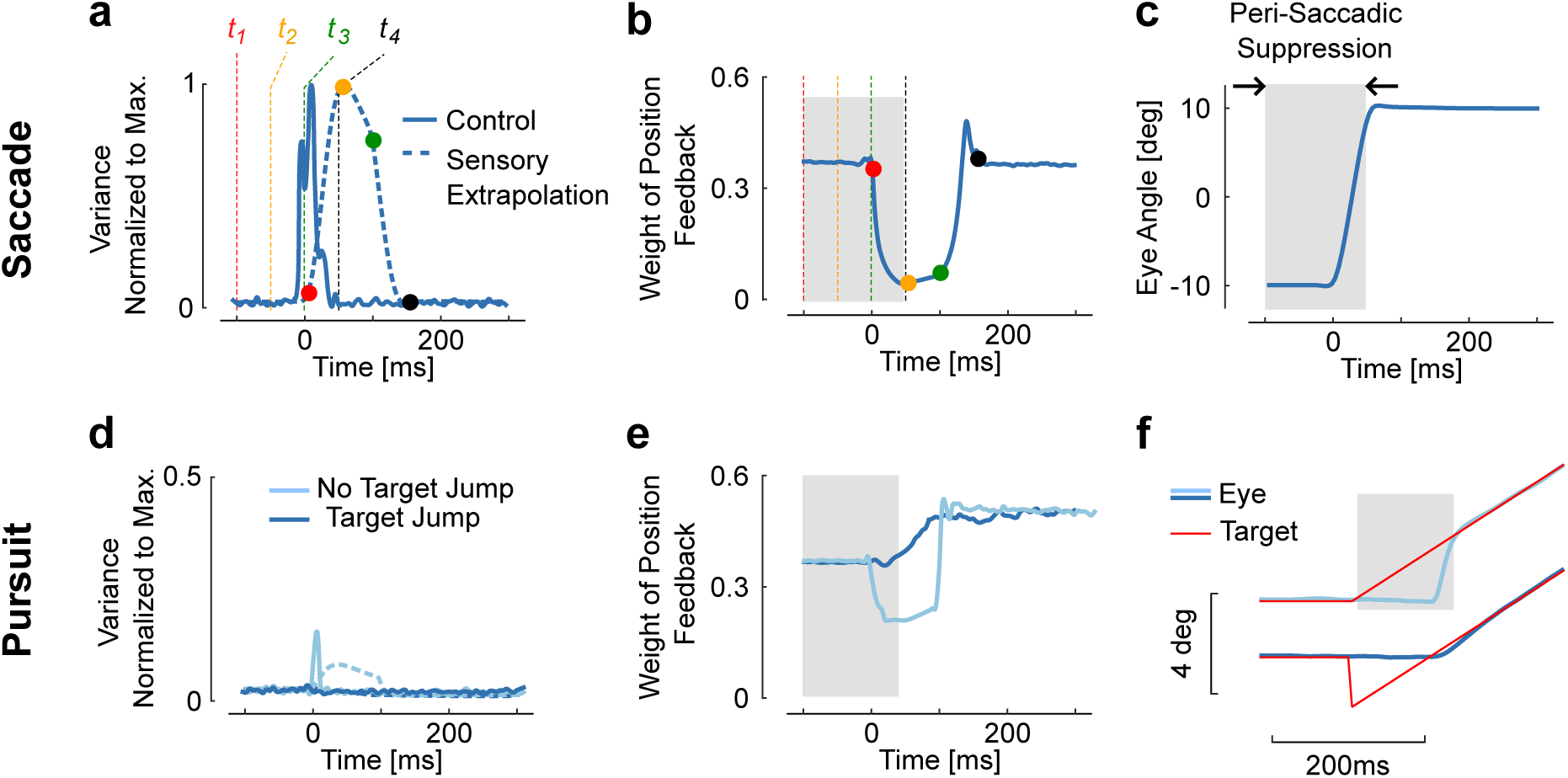
**a.** Variance of the control signal (solid) and of the extrapolation of sensory feedback (dashed). Four times are represented to illustrate how signal-dependent noise impacts the extrapolation of sensory feedback is (*t*_1_ = −100ms, *t*_2_ = −50ms, *t*_3_ = Oms, and *t*_4_ = 50ms). The dots with similar color represent the moment when the information at the corresponding time is available (*t_i_* + 100*ms*). Observe the increase in extrapolation variance associated with stimuli between *t*_1_ and *t*_2_. **b.** Weight of the position feedback for correcting the estimate of the position. This weight is directly taken from the Kalman gain matrix. The reduction in Kalman gain at each selected time point is directly linked to the increase in variance, **c.** Representation of a modeled saccadic eye movement, with the gray area corresponding to the interval of time during which sensory input is given less weight as a result of the extrapolation variance, **d.** Control and extrapolation variance normalized to the maximum values obtained for saccades of 20deg (top traces). **e.** Weight of sensory feedback for the two simulations. Observe that although the catch-up saccade is very small (~2deg), the transient increase in extrapolation variance gives rise to a reduction in weight. **f.** Illustration of the smooth pursuit task with (dark blue) or without (light blue) initial target jump occurring simultaneously with the velocity jump. The absence of target jump evokes a catch-up saccade, which is associated with ta reduction in sensory weight. There is no reduction with the initiation of smooth pursuit.

The time-varying variance induced by control-dependent noise has a direct impact on the importance of visual inputs, through the Kalman gains (Fig. 3b). As the Kalman gain is inversely proportional to the extrapolation variance (see Methods, Eqn. 8), the transient increase in the variance of sensory extrapolation generates a reduction in the weight of sensory feedback about the eye position. And indeed, sensory suppression is seen around the time of simulated saccades (Fig. 3b). The period of suppression is long because high variance period includes the movement time, but also the delay (gray rectangle in Fig. 3c). The model thus predicts that feedback about stimuli from this time window should be given less weight in order to optimally estimate the state of the eye.

We can also see related effects in the simulated pursuit task. The presence of a catch up saccade, even a small one (4deg), is sufficient to evoke a transient increase in extrapolation variance (Fig. 3d-f). This results in a reduction in the weight of sensory feedback. In contrast, when the eye starts moving smoothly (Fig. 3f), there is no catch-up saccade needed and the model predicts no visible change in the weight of sensory feedback. The behavioral finding^39^ that saccades but not smooth pursuit elicit suppression of sensory feedback, and that the suppression scales with the amplitude of the catch-up saccade, directly results from this model.

Thus far, we have described how an efficient state estimator should reduce the weight of sensory feedback around the time of saccade. However, saccadic suppression clearly impacts perception, and it is not obvious how this phenomenon relates to our model. In the introduction we set up two hypotheses. Perception and action could be separate, and in this case the Kalman filter for movement should not affect perception (Fig 1, H1). Alternatively they could share resources, such that a lower Kalman gain for movement should carry through to a lower weight of vision for any estimate, including perceptual ones (Fig 1, H2). If H1 is true, then we should find that both neural and behavioral variables that relate to perception are unaffected by saccades, beyond the limited effect induced by the rapid change of the retinal image ^16^. If H2 is true, on the other hand, we should find suppression not only of behavioral variables but also directly in the neural activities associated with sensorimotor control.

Behaviorally, we can analyze data from perception experiments. Suppression should occur prior to movement onset, reach a maximum close to movement onset (Fig. 4b), and scale with the movement amplitude with relatively invariant timing across amplitudes. Interestingly, this goes even down to the level of microsaccades, inducing partial suppression despite being very small in amplitude ^40^. These known properties of saccadic suppression are in line with the model prediction (Fig. 4b, black): contrasts sensitivity is reduced and visual stimuli such as flashes, gratings, or small displacements are less likely to be accurately perceived ^16,19,20,41–43^. This even happens when the stimuli are chosen so that the eye movement does not change the retinal image, which is compatible with the model (see the simulated contrast reduction of a white stripe, Fig. 4a). Finally, although timing is preserved across amplitudes ^15^, the model predicts that the magnitude of suppression scales with the saccade amplitude as observed experimentally ^44^. This is a direct consequence of signal-dependent noise. Hence, the behavioral data is consistent with Hypothesis 2 where sensorimotor behavior and perception share a common neural resource.

**Figure 4.**
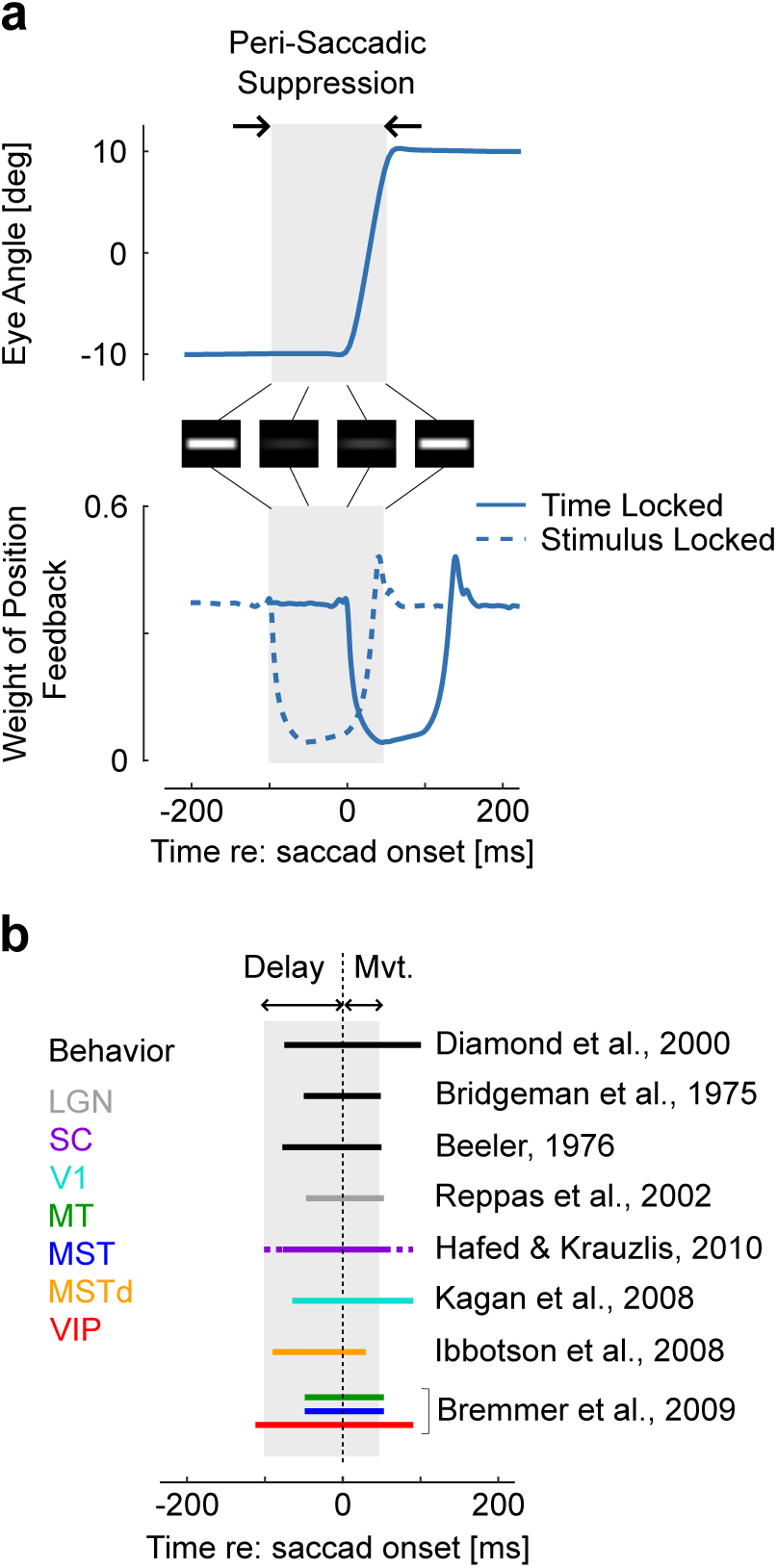
Representation of a simulated 20deg saccade ad dynamic weight of sensory feedback, with the perisaccadic suppression highlighted in gray. These traces are similar as Figure 3a and 3c. The images represent the convolution of a horizontal stripe with a Gaussian kernel with variance proportional to the extrapolation variance to highlight that assigning higher variance may lead to reduced contrast, even when the movement is aligned with the stimulus orientation. Times correspond to the Fig. 3. The decrease in Kalman gain occurs from 0 to 150ms relative to saccade onset (solid trace: time locked), thus the window during which stimuli are suppressed corresponds to −100 to 50ms (dashed trace: stimulus-locked). **b.** Illustration of how the predicted saccadic suppression compares with previously reported suppression from behavioral and neural data. The duration of the perisaccadic suppression in the model is the sum of the temporal delay and of the movement time as represented above with the gray rectangle. Comparisons are approximate as movement time was not the same across all studies. The solid and dashed traces for saccadic suppression in SC indicates the range of onset and offset as given by Hafed and Krauzlis^40^. Other intervals of saccadic suppression were drawn following the authors’ summary or based on visual inspection of the corresponding references.

There is an indirect but interesting parallel between the model's behavior and the selective suppression of stimuli with low frequency contents ^19,20^. The model predicts that sensory feedback about the eye position, but not necessarily velocity, should be reduced. The reason is that the expected signal-dependent noise directly impacts the prediction of the forward model about velocity (which is not the case for position given the current estimates of velocity). Because the predicted and extrapolated velocities are both uncertain, the best strategy is to give them similar weights. This is interesting because changes in velocity typically have higher frequency contents than changes in position, thus a selective suppression of stimuli with low frequency contents may be directly related to the reduction in the weight of feedback about position, but not velocity.

Neurally, we may also expect changes. One obvious way how changes in Kalman gains could be implemented is that neurons could just be driven less strongly when there is more uncertainty. This should predict reduced firing rates around the time of the saccade. Indeed, a large number of experimental studies have found such a neural suppression across the hierarchy of visual areas ^45^. It starts with the lateral geniculate nucleus (LGN)^46^ and the superior colliculus (SC)^40^, via VI^47^, MT, MST, MSTd ^45,48^, all the way up to area VIP ^48^ (Fig 4b, colored). Interestingly, the timing is very similar across brain regions, which emerges naturally from the fact that the loop through the outside world with its delays is the dominating timescale. As predicted by the shared resources model, there is suppression across the entire visual hierarchy.

## Discussion

We have presented a feedback control model that assumes movement dependent-noise and delays, and uses state estimation to optimally control eye movements. It is built on the insight that motor noise is unavoidable and produces sensorimotor uncertainty. It assumes that perception and movement share common resources and hence a common gain on visual input. Behaviorally, it describes the dynamics of both smooth pursuit and saccades. Perceptually, it describes the suppression of sensation around the time of saccades. Neurally, it captures the reduction of neural responses to visual stimuli presented before or during saccades.

A clear limitation of our model is that it is difficult to test experimentally. However the important motivation behind this study was that maintaining a stable visual scene, as commonly assumed, does not explain the phenomenon of saccadic suppression. Indeed, suppression in this case should only occur when the eye moves, and should not be stronger than the moderate loss in performance associated with simulated saccades ^16^. All discarded information beyond movement-related effects would otherwise represent a net loss (up to ~100ms for some brain areas, Fig. 4b). Thus it is clear that saccadic suppression is either very inefficient, or that maintaining a stable visual scene is just not its only purpose. We provide an alternative explanation that captures suppression qualitatively in the context of sensorimotor control. Rather than providing a definite answer to why suppression occurs, we highlight a plausible explanation and expect that it provide an insightful framework for interpreting data about visual processing.

In addition to capturing the major aspects of behavioral and neural suppression, our model explains the previous findings of Watson and colleagues ^41^, who investigated the detection of noisy gratings in human, and found that the best explanation for saccadic suppression was a stimulus-independent reduction in the response gain. This result is a key aspect of saccadic suppression: the retinal images do not become intrinsically noisier; instead it is the visual system that responds less to a given stimulus. Our model also accounts for this result: by reducing the sensory weight in the Kalman gain, the controller becomes less sensitive to sensory information. This is due to the uncertainty induced by the motor commands, which in this case it is clearly independent of the retinal image. The contribution of our model is to show that this stimulus-independent reduction in response gain may be rooted in efficient computations about the state of the eye.

We propose such a mechanism as plausible origin of saccadic suppression, but cannot indicate how the visual system performs this operation at the level of neural circuits. We draw a qualitative link between Kalman filtering and the reduction in sensory weight or neural excitability, and thus this link remains speculative. However, the model does provide hints about what to look for. First, the increase in extrapolation variance clearly results from convolving the motor-dependent noise with the expected eye dynamics over the delay interval. Second, this increase is directly proportional to the integrated motor command. Thus, convolutional networks in the visual system receiving the corollary discharge as input may easily implement a reduction in the gain of neural responses that achieves statistically optimal sensory weighting. Any anatomical or functional similarity between these putative neural operations and neural data may thus provide insight into the circuitry underlying state estimation.

Besides these limitations, we believe that the model is compelling because it is simple. The insight behind our model was to consider both sensory delays and noisy commands together, which are known to play an important role in sensorimotor control ^49,50^. Our approach combines predictive control (based in this case on *finite-spectrum assignment*^51^) and optimal filtering ^52^, to derive a closed-loop controller that handles delays and noise efficiently. In this framework, the weight of feedback must be adjusted not due to movement on its own, but to the uncertainty that results from motor noise and sensory delays. This result, and the diversity of behaviors that the model reproduces, simply fell out of the simplest instance of stochastic optimal control (LQG).

In addition to reproducing saccadic suppression, our model succeeded at the difficult task of controlling fast movements with comparatively long delays, without artificially interrupting the sensory inflow. Indeed, previous models of saccadic control tend to only consider *open-loop* controllers ^49^, or closed-loop control with internal feedback only ^25,53,54^ while it is clear that sensory information remains available and may influence online control^27,28^. Here our model predicts that sensory feedback must be strongly reduced, but not completely suppressed, as observed experimentally ^18^. This is because the Kalman filter achieves an optimal projection in the probabilistic sense, by making the estimation error orthogonal to (or statistically uncorrelated with) the estimated state. Thus, the decrease in the Kalman gain during movement indicates that state information prior to the saccade still carries some information about the current state, and thus can be exploited to derive optimal estimates. The resulting control law (see Methods, Eqn. 10) plays the role of a burst generator, and can be easily inserted as such in more complex models of gaze control.

Perception and control may thus share a common neural substrate in the visual system. The question of why the nervous system shares resources remains open. If perceptual and sensorimotor processes were completely decoupled, we could perform better (Fig. 1, H1). Thus the hypothesis of shared resources must find a good reason to live (Fig. 1, H2). First it is clear that the corollary discharge is critical to monitor the consequences of motor commands. Thus, perception is already conditional upon the ability to integrate extra-retinal signals accurately, both during saccade and pursuit^24,32,40,55^. Second, although perception on its own is suboptimal during saccades (we discard a lot of meaningful information), the shared resources model is clearly cheaper in terms of neural resources. It may thus be globally optimal if the perceptual loss is not more penalizing than the cost of dedicating more neural resources to visual processing in general, considering perceptual and control systems together. Using the same posterior belief for perception and action also ensures self-consistency, in the sense that the same stimulus is not deemed more or less reliable dependent on how we use it. Self-consistency is known to characterize perceptual judgment tasks, where participants make a continuous use of the hypothesis that they previously committed to ^56^. Our model suggests that similar principles may governs the use of posterior beliefs about the state of the eye for perception and control, indicating that these functions emerge from a shared neural substrate. We expect that future work investigate whether this theory is generally applicable to other examples of active sensory suppression associated with voluntary actions such as force generation or reaching^57–59^.

## Methods

### Biomechanical model

We consider a second-order, low-pass filter as a biomechanical model of the oculomotor plant. Based on previous modeling work ^60^ we set the time constants to *τ*_1_ = 224*ms* and *τ*_2_ = 13*ms*. In the sequel, scalars are represented with lower-case characters, vectors with bold lower-case and matrices with capitals. Thus the state-space representation of the continuous-time differential equation representing the eye dynamics was:

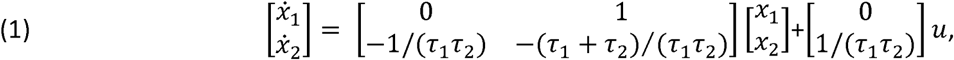

where *x*_1_ is the eye angle, *x*_2_ is the eye velocity, *u* is the command input and the dot operator is the time derivative. The explicit dependency on time was omitted for clarity. This representation takes the form:

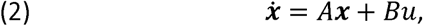

with *x* := [*x*_1_ *x*_2_]^*T*^ representing the state of the system.

The plant model was then augmented with the target position and target velocity, and transformed into discrete time model to include sensorimotor noise. The discrete-time stochastic dynamics and the measurement equations are as follows:

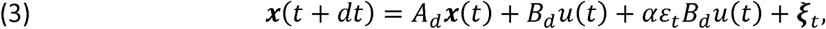

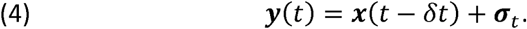

The matrices *A_d_* and *B_d_* form the discrete-time state space representation of the continuous-time system defined in Eqn. 1, which for a discretization step of *dt* corresponds to: *A_d_* = *e^dtA^*, and 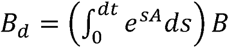 The constant *α* > 0 is a scaling parameter; *ε_t_* and ***ξ***_*t*_ and ***σ***_*t*_ are Gaussian noise disturbances. The multiplicative noise *ε_t_*) is a scalar with zero mean and unit variance, whereas the additive sources of noise are 4-dimensional random disturbances with zero mean and variance set to *∑_ξ,σ_* which will be defined below (recall that the state vector includes that target position and velocity).

The measurement delay was *δ_t_* = 100*ms* in a agreement with measured and modeled latencies of rapid saccadic responses to visual stimuli^61,62^. The subscript *t* for the random noise disturbances was used to remind that these series do not have finite instantaneous variation; but they have finite variance over the discretization interval of *dt*.

### Closed-loop controller

Optimal estimation and control of the stochastic system defined in Eqns. 3 and 4 can be derived in the framework of extended Linear-Quadratic-Gaussian control (LQG), including the effect of control and state-dependent noise ^63^. However this approach is not necessarily well suited for handling sensorimotor delays because it requires system augmentation ^35^, and as a consequence the estimator achieves optimal (probabilistic) projection of the prior estimate onto the delayed state measurement^52^. Since we know that the visual system extrapolates sensory information to compute the present state of the eye or of a moving target^29–31,64^, we were interested to derive an optimal estimator that explicitly extrapolates sensory signals, captured in ***y***(*t*), over the interval *δ_t_* (see also Fig. 2). The key aspect of this estimator design is that, by taking into account the control function *u*(*s*), *t* – *δt* ≤ *s* ≤ *t*, the variance of the extrapolated sensory signal is dynamically adjusted as a function of the control-dependent noise (3^rd^ term of Eqn. 3).

More precisely, we assume that neural processing of sensory signals consists in computing an estimate of the present state of the eye given the delayed sensory signals as follows:

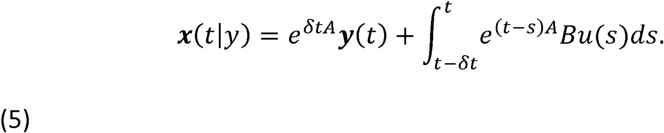

Using the notation *M*(*t*):= *e^tA^*, it is easy to observe that the extrapolation error (Δ_*t*_) follows a Gaussian distribution defined as follows:

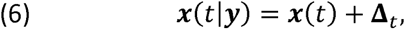

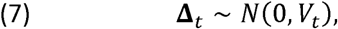

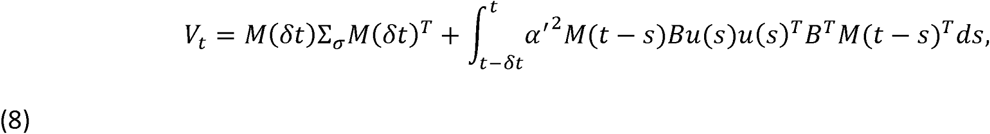

where *α′* = *α*(*dt*)^−1/2^ was defined in agreement with the unit-variance Brownian noise disturbance considered for the stochastic differential equation.

With these definitions, we can derive an adaptive estimator based on standard Kalman filtering using the extrapolated state (Eqn. 8) instead of the available state measurement (Eqn. 4). The state estimate is computed in two steps as follows:

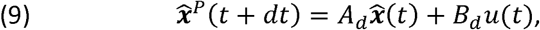

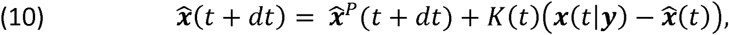

and the Kalman gain, *k*(*t*), well as the covariance of the estimated state are updated iteratively following standard procedures^52^.

Observe that the separation principle does not hold because the variances of the one-step prediction and of **Δ**_*t*_ both depend on *u_t_*. Thus our approach is valid under the assumption that the control must not be jointly optimized with the state estimator. Instead of optimizing iteratively the controller and the state estimator as in the extended LQG framework ^63,65^, we computed the controller independently based on the heuristic assumption that the separation principle applied, and then optimized the state estimator defined in Eqns. 9–10 by taking into account the effect of control-dependent noise explicitly (Eqns. 6–8). The controller was thus obtained by solving the LQG control problem while ignoring the multiplicative noise in Equation 3 as follows:

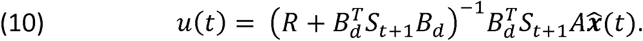

In Eqn. 10, *R* represents the cost of motor commands, and the matrices *S_t_* are computed offline following standard procedures ^63,66^.

We developed this approach to include the extrapolation of sensory data while considering the control over the delay interval explicitly. Using feedback control based on a predicted state is known as *finite spectrum assignment* (FSA), which is germane to a Smith predictor in that it aims at removing the delay from the feedback loop^51^. Here, FSA was chosen to reconstruct the predicted state (instead of the system output as for the Smith predictor), allowing the use of position and velocity estimates in the control law (Eqn. 10).

### Numerical Simulations

The only free parameters are *α* (the scaling of the signal dependent noise), the covariance matrices of ***ξ***_*t*_ and ***σ***_*t*_ (respectively ∑_*ξ*_ and ∑_*σ*_), and the cost-function used for control. We used the following values: the constant *α* was set to 0.08, ∑_*ξ*_ was 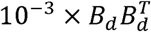and ∑_*σ*_ was 10^−6^ times the identity matrix of appropriate dimension. These parameters were manually adjusted so that when adding the signal-dependent term to the variance of the extrapolation error (Eqn. 8), the Kalman gains converged to steady-state values and the variances of the extrapolated state and of the motor noise were comparable during fixation. It is clear that changing the noise parameters may influence the results qualitatively. However the key feature of the adaptive estimator is that the extrapolation variance increases monotonically with the square of the motor command, which is why the extrapolated measurement is dynamically reduced during movement. This aspect does not depend on the different noise parameters.

The cost parameters were adjusted to generate simulated saccades compatible with typical recordings of eye movements in humans, and these parameters do not impact the results qualitatively. For saccadic movements, we simulated two fixation periods at the initial 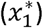 and final 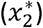 targets during which the cost of position error was 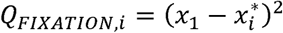 The two fixation periods were separated by the movement time, which was a 50ms window during which the eye was free to move without any penalty on the state vector. For the smooth movements in response to velocity jumps, we simulated a fixation to the target and changed the target state during a simulation run. The cost of motor commands in all cases was *Ru*(*t*)^2^, with *R* := 0.01. Finally we used a discretization step of 5ms.

One difficulty is that the extrapolation requires that all state variables, including the target, be observed independently (Eqn. 4). This is not fully compatible with the visual system, because there is no measurement of the target state independent of the state of the eye. This limitation could be overcome by considering another observer that reconstructs the state vector prior to extrapolating the sensory feedback. Here, instead of considering such additional observer, we injected similar amounts of signal-dependent noise in the sensory feedback about the state of the eye as about the state of the target. This procedure was chosen for simplicity and captures the intuitive idea that if the eye position is very noisy, then information about the target location logically shares the same uncertainty.

## Acknowledgements

The authors would like to thank G. Blohm for comments on an earlier version of this manuscript. FC is supported by a grant from F.R.S.-FNRS (grand number : 1.B.087.15F *Chargé de Recherches*, Belgium) and by grants of the Interuniversity Attraction Poles of the Belgian Science Policy (IPA Network on Dynamical Systems, Control and Optimization, DYSCO).

